# Regular parental exercise before mating influences offspring lower adiposity associated to hypothalamic neurodevelopmental changes

**DOI:** 10.64898/2026.01.10.698777

**Authors:** Celia Muñoz, Carla Canal, Joel Vizueta, Paula Sanchis

## Abstract

Physical inactivity is highly prevalent worldwide and affects not only individual health but also the health of future generations. However, the impact of parental physical activity, limited to the pre-mating period, on offspring body weight and composition remains poorly understood. Using a voluntary wheel running approach in mice, we uncovered that post-weaning offspring body weight and composition changes are modulated by the combined effects of pre-mating parental exercise and parental age. Notably, during lactation, pre-mating parental exercise reduced offspring visceral and subcutaneous adiposity, shortened tibia length in female offspring, and influenced offspring transcriptomic profiles of the hypothalamus, the central region regulating body weight and energy balance. These results highlight that regular pre-mating parental exercise may induce offspring neurodevelopmental changes. Although pre-mating exercise minimally impacted the expression of lactation-related genes in maternal subcutaneous fat, as well as breastmilk nutritional composition and miRNA content, these modest miRNA changes may nonetheless influence offspring hypothalamic regulation. Together, these data provide a comprehensive understanding of how parental age and pre-mating exercise impact post-weaning offspring body weight and composition and offer deeper insights into how regular pre-mating parental exercise influences offspring physiology during lactation.

## INTRODUCTION

Physical inactivity has a range of health implications not only for the individual [1], but also for their progeny [2]. In fact, it has been observed that children of active parents are less likely to be overweight than those from inactive parents [3]. Furthermore, the high prevalence worldwide of inactivity, together with the delay in the timing of conception of the first child[4] is resulting in an elevated risk of a child being born from inactive and old parents [5–7].

In rodents, maternal exercise before and during pregnancy improves offspring glucose metabolism and ameliorates the effects of maternal high-fat-diet on metabolic health in both female and male adult offspring [8–13]. Moreover, maternal exercise before and during pregnancy promotes interscapular brown adipose tissue activity and browning of inguinal white adipose tissue in the offspring [14], and increases exercise-induced placental superoxide dismutase 3, which induces hepatic epigenetic changes in the offspring [8–13]. Moreover, maternal exercise before and during pregnancy protects adult mouse offspring from diet-induced obesity through circulating and peripheral changes in skeletal muscle and adipose tissue [15,16]. Similarly, paternal exercise also improves offspring metabolic and cognitive behavior health and mitigates the detrimental effects of a paternal high-fat diet on offspring health, via epigenetic changes in the sperm [17–20]. Interestingly, the combination of maternal and paternal exercise induces summative effects on offspring glucose tolerance associated with endocrinal changes on pancreas [21], and the grandparental exercise also impacts metabolic health of second-generation offspring [22,23]. Parental age at childbirth might be another factor that may contribute to the recent obesity pandemic, as it is associated with pregnancy, perinatal, and offspring metabolic complications [24–26]. Given that exercise is not always feasible during pregnancy, it remains unclear whether regular pre-mating exercise of dams and sires is sufficient to modulate offspring body composition and energy balance both baseline and metabolically challenged conditions, and whether these effects are influenced by parental age.

It has long been recognized that physical activity influences energy balance through a complex interaction between energy intake and energy expenditure [27]. The brain, and specifically, the hypothalamus plays a central role in the regulation of whole-body energy balance by controlling both food intake and energy expenditure [28,29]. Previous studies have shown that maternal obesity influences the postnatal transcriptomic landscape of the offspring hypothalamus [30], which highlights the impact of parental lifestyle on hypothalamic profile. At present, the direct effect of regular exercise on the transcriptomic signature in the offspring hypothalami is unknown. Therefore, we have conducted a comprehensive physiological study of body weight and composition in offspring, under both baseline and metabolic challenge conditions, to examine the effects of regular pre-mating parental exercise and parental age. Furthermore, we examined how pre-mating parental exercise affects breast milk and the offspring hypothalamus during lactation, providing insights into hypothalamic alterations mediated by parental exercise before mating.

## RESULTS

### Running increased food intake in mice and yielded sex- and age-dependent dimorphic changes in body composition

We first examined whether regular exercise exerts similar effects on body weight and composition in male and female mice of different ages, which could differentially influence their progeny. We monitored single-housed *ad libitum*-chow-fed male and female young and old mice assigned by baseline traits to active (functional wheel) or sedentary (blocked wheel) groups for 5 weeks. In males, the voluntary wheel running increased over time in both young and old mice triggering an increase in energy expenditure, food intake, and fat loss (Fig. 1A-D and S1A-B) as previously seen [31]. Young groups gained lean mass due to somatic growth whereas old groups lost lean mass, which resulted in accentuated body weight loss in old runners (Fig. 1E-F). Despite these changes, no significant differences in estimated energy balance were observed between groups (Fig. S1C). In females, running distance also increased over the time causing an increased energy expenditure and food intake (Fig. 1G-I and S1D-E). Old runners lost fat mass, whereas young runners were resistant to it (Fig. 1J). Given that lean mass increased in both running groups, but more markedly in the younger group, this resulted in similar body weight gain to sedentary mice (Fig. 1J-L). Similar to males, running resulted in no energy balance differences between age-matched groups (Fig. S1F). Collectively, this study reveals age-dependent sex differences in the body composition in response to exercise.

**Figure 1.**
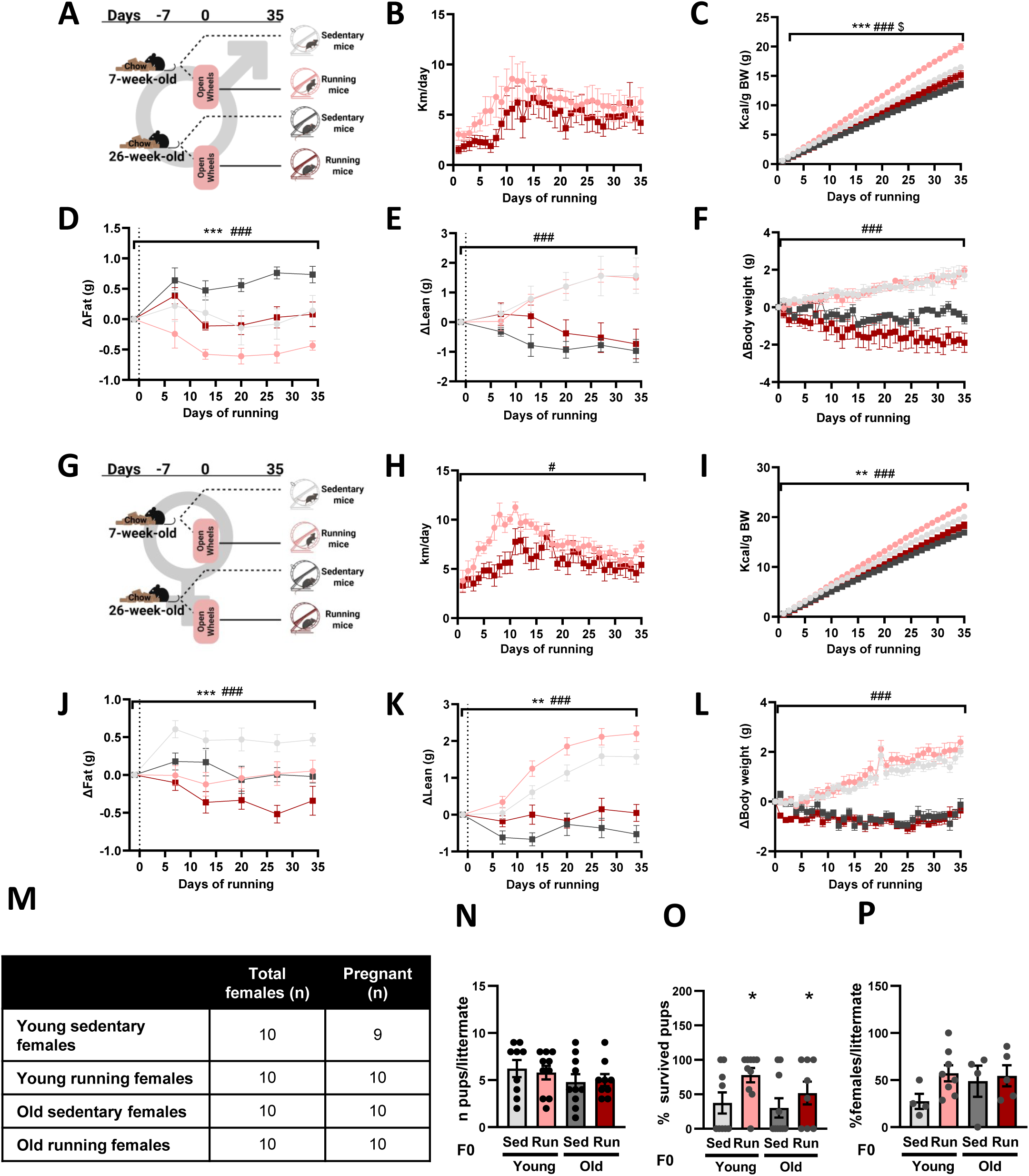
Regular running impacts on body composition in a sex- and age-dependent manner. **A-F**) Regular exercise in young and old males. **A**, Schematic. **B**, Distance. **C**, Food intake. **D**, Fat mass changes. **E**, Lean mass changes. **F**, Body weight changes. **G-L**) Regular exercise in young and old females. **G**, Schematic. **H**, Distance. **I**, Food intake. **J**, Fat mass changes. **K**, Lean mass changes. **L**, Body weight changes. **M-P**) Pregnancy outcomes in response to parental age and pre-mating parental exercise. **M**, Table of pregnant females. **N**, Pups per littermate. **O**, Survived pups. **P**, Females pup rate. One-way repeated-measures ANOVA was used to assess running (*) in **B** and **H**, and two-way repeated-measures ANOVA was used to assess running (*), age (#) and its interaction ($) in **C**-**F** and **I**-**L**, but if Mauchly’s sphericity test and/or Shapiro-Wilk’s Normality test was significant, then repeated-measures Generalized Estimating Equations (GEE) was conducted. Two-way ANOVA was used in **N**-**P**, but if Shapiro-Wilk’s Normality test and/or Levene’s test for homogeneity of variances was significant, Generalized Linear Model (GzLM) was used. Time-course data are presented as mean ± SEM, and bar graphs show mean ± SEM with individual mice represented as dots. *P≤0.05, **P≤0.01, ***P≤0.001. **A** and **G** were created using Biorender.

Given the extensive evidence that parental exercise before and during conception impacts offspring metabolism, we next evaluated whether regular running exclusively during the pre-mating period affects pregnancy outcomes, and whether these effects are influenced by parental age. Most females were pregnant and gained similar body weight regardless of age or regular pre-mating exercise (Fig. 1M; Fig. S1G-H). However, after giving birth, older females weighed more than younger females, as they were before pregnancy, with pre-mating exercised dams weighing more than sedentary ones (Fig. S1G-H).

At weaning, body weight was higher in older mothers due to their greater fat mass, with no differences in lean mass between groups, likely reflecting the reduced lean mass gain in the pre-mating exercise dams (Fig. S1I-L). Although paternal age or regular pre-mating parental exercise do not impact on the number of pups per litter, pre-mating exercise increased the pup survival rate and trended toward increasing the proportion of female pups per litter between younger groups (p=0.052; Fig. 1N-P), in line with the female-biased litters previously reported following parental exercise [32].

### Post-weaning offspring body weight and composition changes are influenced by the interaction of regular pre-mating exercise, parental age, and offspring sex

To evaluate the effects of regular pre-mating parental exercise and parental age on offspring body weight and composition, we selected only one male and one female from litters containing more than six pups, in order to minimize littermate effects [33]. While parental age increased body weight at postnatal day 14 and 21 (Fig. 2A-C), pre-mating parental exercise increased body weight gain during the late lactation period (Fig. 2D). After weaning, both female and male offspring from older parents exhibited increased fat mass during youth compared to those from young parents (Fig. 2E). However, fat mass gain during this period was more pronounced in females, with a significant interaction among parental age, parental exercise, and offspring sex, suggesting that fat mass gain may be modulated in a sex-dependent manner by both parental age and exercise (Fig. 2F). Lean mass, and consequently body weight, was higher in male offspring during youth, predominantly driven by increased lean mass gain in those born to young exercised and old sedentary parents (Fig. 2G-J). These results highlight that offspring body composition and weight changes are influenced by regular pre-mating parental exercise and parental age in an offspring sex-dependent manner.

**Figure 2.**
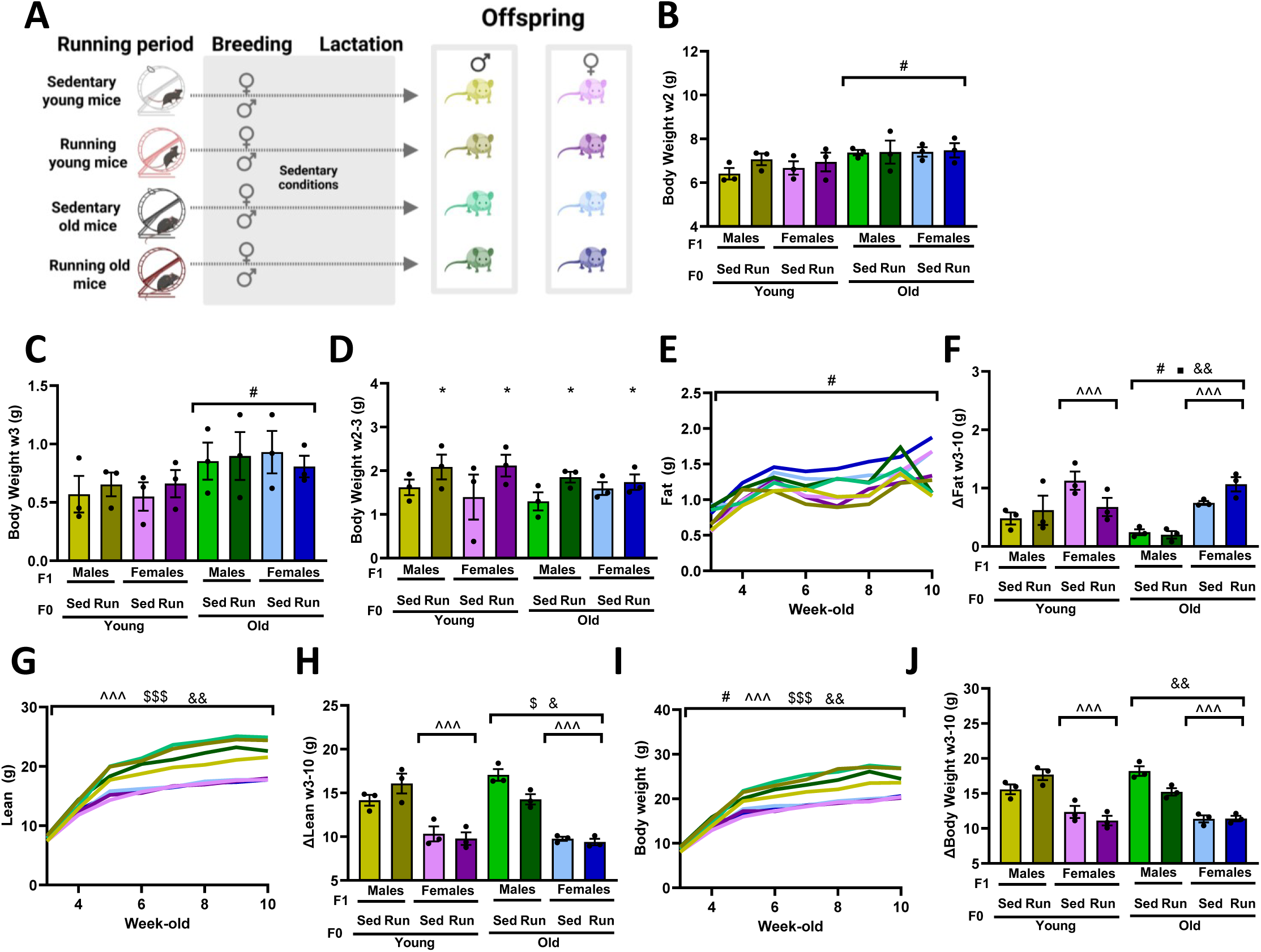
Post-weaning offspring body weight and composition changes are influenced by the interaction of regular pre-mating exercise, parental age, and offspring sex. **A**, Schematic. **B**, Body weight at 2-week-old. **C**, Body weight at 3-week-old. **D**, Body weight changes at weaning. **E**, Fat mass. **F**, Fat mass changes. **G**, Lean mass. **H**, Lean mass changes. **I**, Body weight after weaning. **J**, Body weight changes. Three-way ANOVA was used for evaluating the effects of parental running (*), parental age (#), offspring sex (^) and its interaction ($ for parental running*parental age, ¤ for parental running*offspring sex, • for parental age*offspring sex, and & for triple interaction) in **B**-**D**, **F**, **H** and **J** but if normality test and/or homogeneity of variance were violated, GzLM was used. Two-way repeated-measures ANOVA was used in **E**, **G** and **I**, but if sphericity and/or normality assumptions were significant, then repeated-measures GEE was conducted. Time-course data are presented as mean ± SEM, and bar graphs show mean ± SEM with individual mice represented as dots. *P≤0.05, **P≤0.01, ***P≤0.001. **A** was created using Biorender.

### Changes in offspring body weight and composition in response to high-fat diet and exercise during youth are modestly dependent on parental age and pre-mating exercise

Next, 1 female from each litter with more than six pups was challenged with a high-fat diet (HFD; Fig. 3A). Parental age, but not pre-mating exercise, influence body composition and weight after HFD exposure, mainly after 2 and 6 days of exposure (Fig. 3B-D), and downregulated expression of *Lrrc8aR*, which is related to adipogenesis [34], in the subcutaneous fat at 2-week of HFD (Fig. S2A-B). However, neither parental age nor pre-mating parental exercise influenced the percentage of regional adiposity (Fig. 3E) or the expression of genes in visceral fat, hypothalamus, or other fat-related genes such as *Leptin*, thermogenesis-related genes like *Adrb3,* or triglyceride mobilization genes such as Lipase E (*Lipe*) in subcutaneous fat (Fig. 3B-D). The tibia length, and thus, overall linear growth of female offspring also remained unchanged by parental age and pre-mating exercise at youth (Fig. 3F).

**Figure 3.**
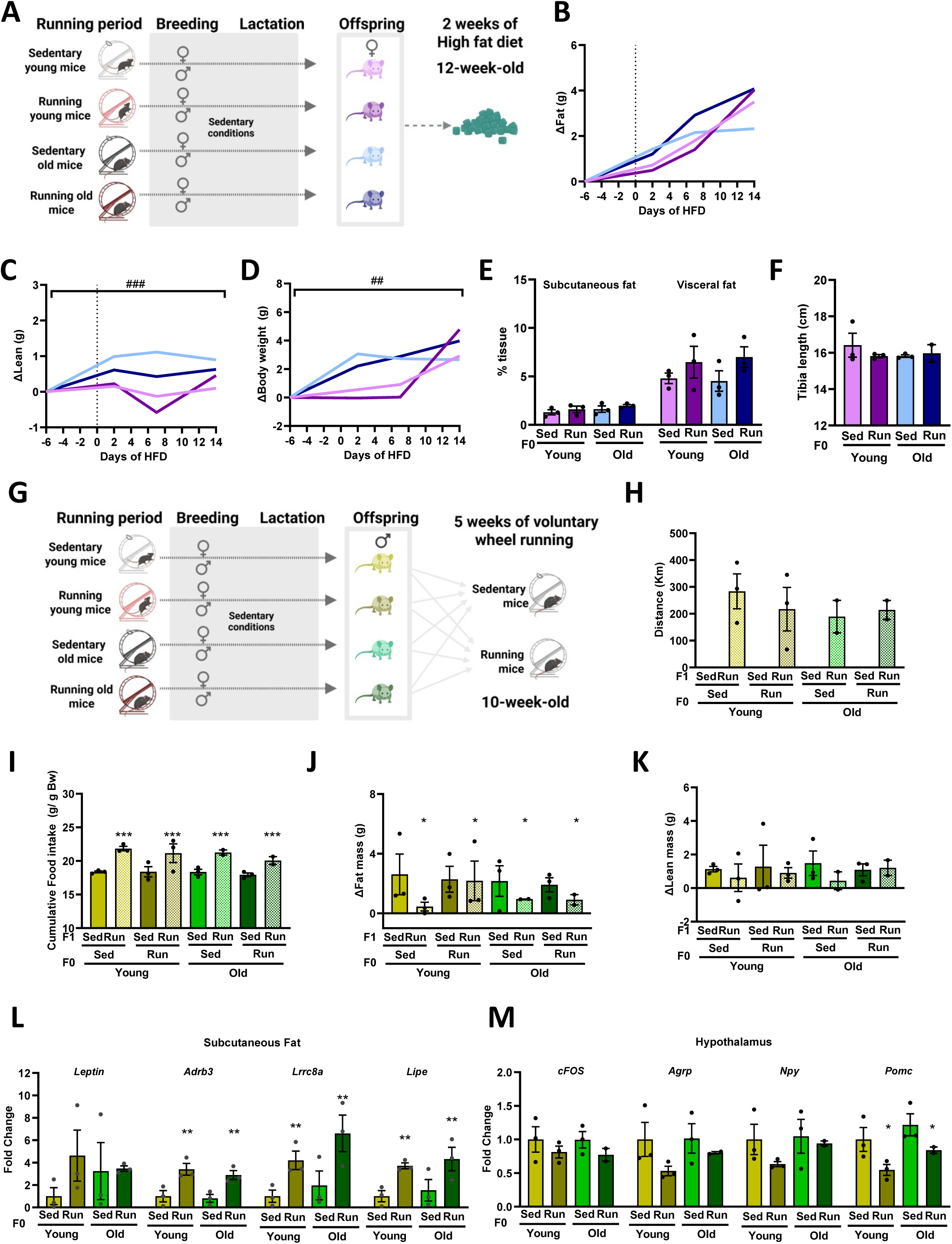
Offspring body composition changes in response to a high-fat diet and exercise during youth are modestly influenced by parental age and regular pre-mating exercise. **A-F**) Female offspring were fed a HFD for 14 days. **A**, Schematic. **B**, Fat mass changes. **C**, Lean mass changes. **D**, Body weight changes. **E**, Tissues. **F**, Tibia length. **G**-**M**) Male offspring were undergo voluntary wheel running protocol. **G**, Schematic. **H**, Distance. **I**, Food intake. **J**, Fat mass changes. **K**, Lean mass changes. **L**, Gene expression changes in the subcutaneous fat from sedentary offspring. **M**, Gene expression changes in the hypothalamus from sedentary offspring. Two-way repeated-measures ANOVA was used to assess running, age (#) and its interaction in **B**-**D**, but if sphericity and/or normality were significant, then repeated-measures GEE was conducted. Two-way ANOVA was used for evaluating the effects of parental running (*), parental age and its interaction in **E**, **F**, **H**, **L** and **M**, and Three-way ANOVA in **I**-**K**, but in cases of normality and/or homogeneity of variance violations, GzLM was used. Time-course data are presented as mean ± SEM, and bar graphs show mean ± SEM with individual mice represented as dots. *P≤0.05, **P≤0.01, ***P≤0.001. **A** and **G** were created using Biorender.

Given the null effect of parental exercise on HFD feeding in females, 1 male from each litter with more than six pups was given access to a running wheel, while another male from the same litter had wheel access blocked (Fig. 3G). No differences in running distance were observed due to parental age or pre-mating exercise (Fig. 3H). As expected, energy expenditure and food intake increased whereas fat mass decreased in response to running (Fig. 3I-K and Fig. S2E-2I), as previously observed, resulting in a total energy balance reduced in running mice (Fig. S2J). However, the parental age and regular pre-mating exercise interacted to modulate the lean mass and body weight during running (Fig. S2F-H). No changes were observed regarding tibia length (Fig. 3K). These findings suggest that the exercise-induced increases in food intake, energy expenditure, and fat loss observed in offspring runners are conserved mechanisms of running, independent of parental pre-mating exercise or age.

Next, we focused on elucidating the effects of parental age and exercise on sedentary offspring, given the lack of effects observed during the running period. Although no effect of parental age and exercise on regional adiposity or liver mass was observed (Fig. S2L), parental exercise upregulated *Adrb3*, *Lrrc8aR*, and *Lipe* genes in subcutaneous fat (Fig. 3L). However, this effect was not shown in the visceral fat (Fig. S2M), suggesting a specific role of pre-mating exercise on offspring subcutaneous fat. In the offspring hypothalamus, parental exercise downregulated *Pomc* (Fig. 3M), which encodes anorexigenic peptides, but did not affect other genes related to feeding behavior (Fig. S2N). Together, these data reveal that, although parental exercise was conducted pre-mating, it can induce changes in the expression of specific genes in offspring tissues during youth.

### Regular exercise modulates locomotion and anxiety-like behavior in response to new environments during the running period, but does not influence maternal nursing behavior after running period

Considering that weight gain of offspring differed between offspring of pre-mating exercised and sedentary parents over this transitional feeding phase (Fig. 2C), which corresponds to the period when mice begin to eat solid food [35], a second cohort of mice was bred following the same group strategy as before (Fig. 4A) to analyze the effect of parental exercise during lactation.

**Figure 4.**
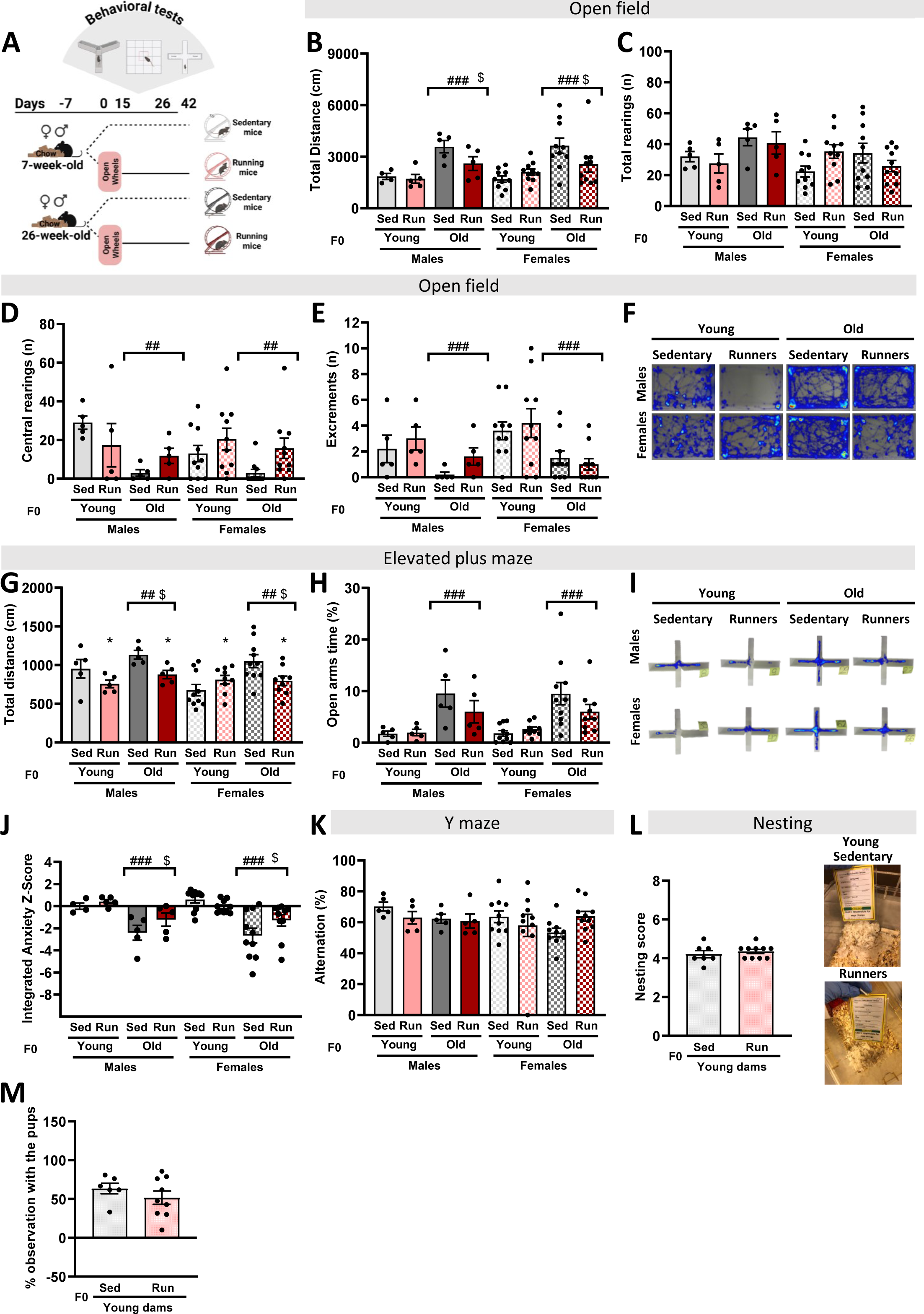
Locomotion and anxiety are influenced by age and regular exercise. **A, Schematic. B-F**) Open field (OF) results. **B**, OF total distance. **C**, OF total rearings. **D**, OF central rearings. **E**, OF excrements. **F**, OF representative images. **G**-**I**) Elevated plus maze (EPM) results. **G**, EPM total distance. **H**, EPM time in open arms. **I**, EPM representative images. **J**, Integrated Anxiety Z-score. **K**, Alternations in Y-maze. **L**, Nesting score in pregnancy young females. **M**, Duration of maternal interaction with offspring. Three-way ANOVA was used for evaluating the effects of parental running (*), parental age (#), parental sex (^) and its interaction ($ for parental running*parental age, ¤ for parental running* parental sex, • for parental age* parental sex, and & for triple interaction) in **B**-**E**, and **G**-**H**, **J**-**K** but if Shapiro-Wilk’s Normality test and/or Levene’s test for homogeneity of variances was significant, GzLM s was used. Unpaired t-test was used in **L**-**M**. Bar graphs show mean ± SEM with individual mice represented as dots. *P≤0.05, **P≤0.01, ***P≤0.001. **A** was created using Biorender.

This second cohort also displayed age-dependent sex differences in the body composition in response of exercise (Fig. S3A-E). During running period, we analyzed parental behavior in response to exercise and age (Fig. 4A). Aging increased total distance traveled, decreased central rearings and fecal excrements in the open field (OF; Fig. 4B-F), and increased both total distance traveled, and time spent in the open arms of the elevated plus maze (EPM; Fig. 4G-I), collectively indicating increased locomotion activity and reduced anxiety-like behavior in old mice. However, exercise interacted with age reducing the total distance in older females in OF (Fig 4B) and decreased total distance in the EPM in running mice (Fig. 4G). To gain further clarity from our behavioral findings, we calculated an integrated anxiety z-score (Fig. 4J), which provides a general assessment of anxiety-like behavior [36,37]. The integrated anxiety z-score, derived from the OF and EPM, revealed that exercise increased anxiety of old females to young levels (Fig. 4J). The Y-maze, which was used to evaluate short-term spatial working memory, was not affected by either regular running or age (Fig. 4K). Together, these findings underscore that regular exercise influences anxiety-like behavior in an age-dependent manner, which may be partly explained by changes in locomotor activity.

Only 5 out of 20 older mothers gave birth, compared to 17 out of 20 younger mothers, with 4 out of 5 successful pregnancies occurring in the exercise group (Fig. S3F). Given the limited number of littermate in old mothers (Fig. S3G-I), we focused exclusively on offspring from young parents. To evaluate whether pre-mating exercise and parental age could affect maternal behavior traits and, in turn, influence early development and offspring survival [38,39], we evaluated the nesting and maternal behavior. Parental pre-mating exercise did not significantly affect nesting behavior on day 17 post-breeding (Fig. 4L), nor did it enhance the time that dams spent with their pups on day 7 postpartum (Fig. 4M), suggesting that pre-mating exercise does not interfere with maternal nursing behavior.

### Pre-mating parental exercise reduces offspring visceral and subcutaneous regional adiposity during lactation and modulates offspring hypothalamic signature

Based on previous considerations [33], we standardized the litter size neonatally to number of 6 pups per littermate on day 2 to avoid differential results due to litter size. Considering that offspring utilize milk most efficiently for weight gain before day 7, and that peak milk production occurs between days 10 and 16 of lactation [40], we analyzed 1 male and 1 female pups on postnatal day 9 per littermate based on mean body wight per littermate and sex. Although no significant differences in body weight were observed between offspring from sedentary and running mice (Fig. 5A-B), offspring from pre-mating exercised parents displayed less visceral and subcutaneous fat than those from sedentary parents (Fig. 5C-D). Although females displayed more brown fat, there were no differences regarding regular pre-mating exercise (Fig. 5E). A trend to shorten the tibia length was observed regarding parental exercise (p=0.072) during lactation, which was more pronounced in females (t-test; p=0.005; Fig. 5F). These data reveal that pre-mating exercise influences offspring physiology during exclusive milk-feeding period.

**Figure 5.**
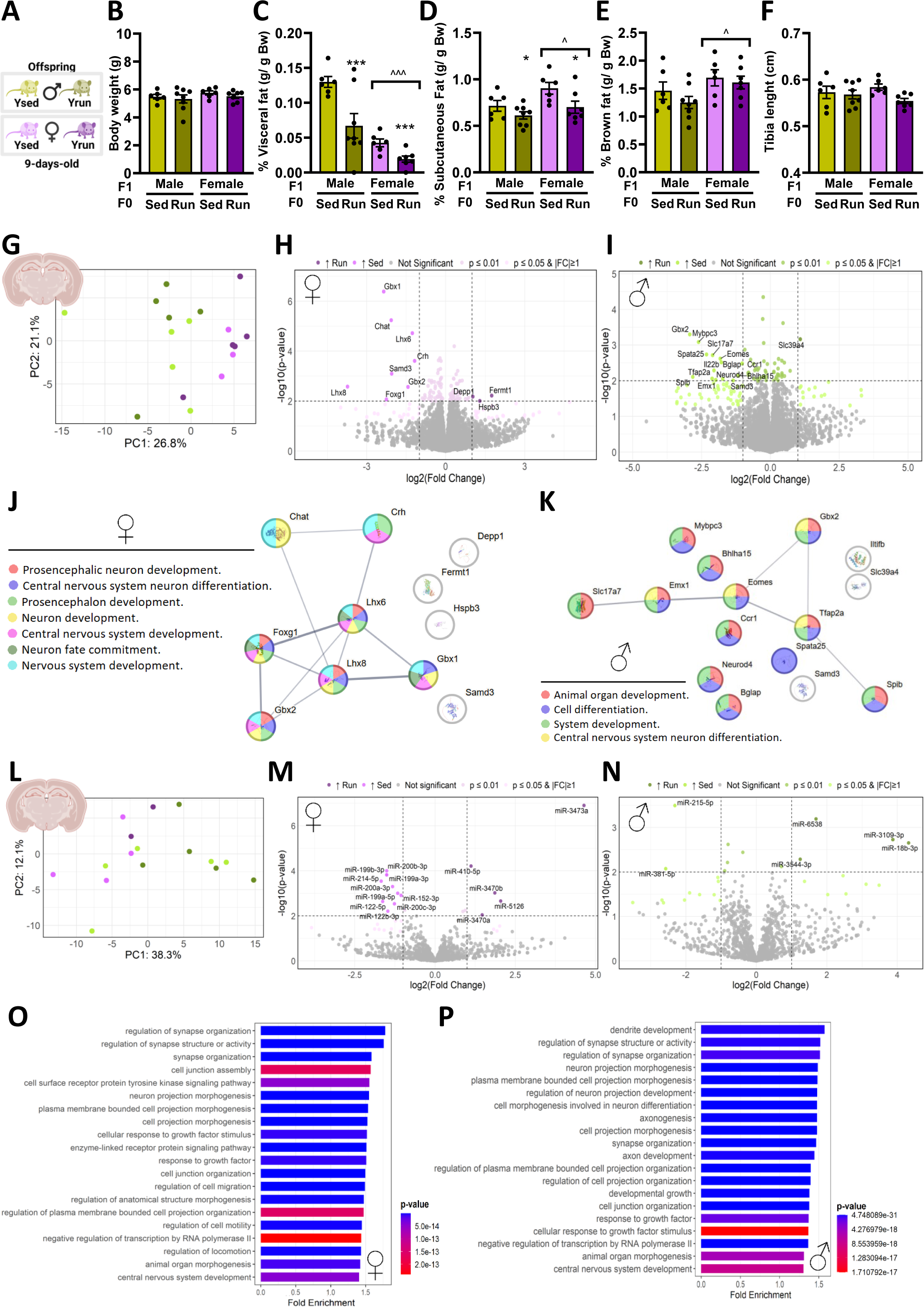
Offspring hypothalamic adaptations of sedentary and pre-mating exercised dams. **A-F**) In vivo results. **A**, Schematic. **B**, Body weight. **C**, Visceral fat. **D**, Subcutaneous fat. **E**, Brown fat. **F**, Tibia length. Two-way ANOVA was used for evaluating the effects of parental running (*), offspring sex (^) and its interaction ($) in **B**-**F**, but if Shapiro-Wilk’s Normality test and/or Levene’s test for homogeneity of variances was significant, GzLM s was used. Bar graphs show mean ± SEM with individual mice represented as dots. *P≤0.05, **P≤0.01, ***P≤0.001. **G-K**) Hypothalamic transcriptomic profiles of male and female offspring in response to pre-mating parental exercise. **G**, PCA of offspring transcriptomic profile. **H**, Volcano plots of DE genes in female offspring from sedentary versus pre-mating exercised dams. **I**, Volcano plots of DE genes in male offspring from sedentary versus pre-mating exercised dams. **J**, STRING analysis of DE genes from female offspring. **K**, STRING analysis of DE genes from male offspring. **L**, PCA of offspring miRNA profile. **M**, Volcano plots of significantly different miRNAs in female offspring from sedentary versus pre-mating exercised dams. **N**, Volcano plots of significantly different miRNAs in female offspring from sedentary versus pre-mating exercised dams. **O**, GO biological process enrichments of miRNAs from female offspring. **P**, GO biological process enrichments of miRNAs from female offspring. **A**, **G** and **L** were created using Biorender.

Given the similarity in the expression of genes related to lipid metabolism, adipogenesis, thermogenesis, and cell proliferation in subcutaneous and brown adipose tissue of male and female offspring in response to parental exercise or age (Fig. S4A-C), we next evaluated the hypothalamic transcriptomic signature of male and female offspring during lactation in response to parental exercise. PCA showed a maximum clustering due to sex but minimal clustering regarding pre-mating parental exercise (Fig. 5G). Similarly, the distribution of cell proportions estimated from transcriptomic deconvolution revealed sex-specific differences in dividing and immune cells, while no effect of regular pre-mating exercise was detected (Fig. S4D). Using a threshold of |fold change| > 2 and p < 0.05, we identified 15 genes (1 upregulated and 14 downregulated; Fig. 5H) and 11 genes (3 upregulated and 8 downregulated; Fig. 5I) differentially expressed (DE) genes in hypothalamus from female and male offspring mice, respectively. Notably, 2 genes (Samd3 & Gbx2, both downregulated) were commonly dysregulated in both comparisons. Most of the downregulated genes by pre-mating parental exercise were related to neurodevelopmental processes (Fig. 5J-K and S4E-F).

MiRNAs are emerging as an important epigenetic mechanism that regulates gene expression not only at the translational level but also through DNA methylation [41]. Therefore, we next evaluated the hypothalamic miRNAs content in response to pre-mating parental exercise. PCA also showed a clustering due to sex, minimal clustering regarding pre-mating parental exercise (Fig. 5L). Using a threshold of |fold change| > 2 and p < 0.05, we identified 15 (5 upregulated and 10 downregulated; Fig. 5M) and 6 (4 upregulated and 2 downregulated; Fig. 5N) differentially expressed miRNAs in hypothalamus from female and male offspring mice, respectively. Although no miRNAs were commonly dysregulated in both comparisons, those dysregulated were also related to neurodevelopmental processes (Fig. 5O-P). Together, these data suggest that pre-mating parental exercise may influence offspring neurodevelopment during lactation.

### Pre-mating parental exercise may influence hypothalamic regulatory networks in offspring

Considering that mice are exclusively breastfed on day 9 [40], we speculated that milk composition could be altered in response to pre-mating exercise. Thus, we evaluated milk nutritional composition on day 9. We did not observe differences in free fatty acids or proteins (Fig. 6A–B). However, there was a trend toward decreased milk carbohydrate concentration in response to pre-mating exercise in dams (Fig. 6C). Next, we analyzed the subcutaneous mammary fat on the same day and observed a significant downregulation of *Lipe*, along with a trend toward reduced expression of *Leptin* and *Adiponectin* (Fig. 6D). Exercise is known to modulate adipose tissue and decrease circulating leptin levels in parallel with reductions in total fat mass [42]. In this context, the downregulation of *Adiponectin* and *Lipe*, both key regulators of lipid metabolism and lipolysis [43], suggests diminished adipocyte mobilization, likely due to decreased, maternal fat reserves in runners (Fig. 6D). In contrast, we observe no differences in the expression of genes related to adipogenesis such as *fatty acid synthase* (*Fasn*), *Fatty acid desaturase* 1 (*Fads1*) and *Lrrc8aR*; thermogenesis evaluated by *Adrb3* expression; the cell proliferation marker *Ki67*; *Signal Transducer and Activator of Transcription 5* (*STAT5*), a key regulator of mammary gland development, remodeling, and milk production [44], or other milk and lactation support proteins such as *Casein Beta* (*Csn2*) , *Prolactin*, *Milk Fat Globule-EGF Factor 8* (*Mfge8*) and *Lactalbumin Alpha* (*Lalba*; Fig. S5A-B). Together, these results highlight that pre-mating exercise had no impact on milk composition and modestly influenced gene expression in maternal subcutaneous fat.

**Figure 6.**
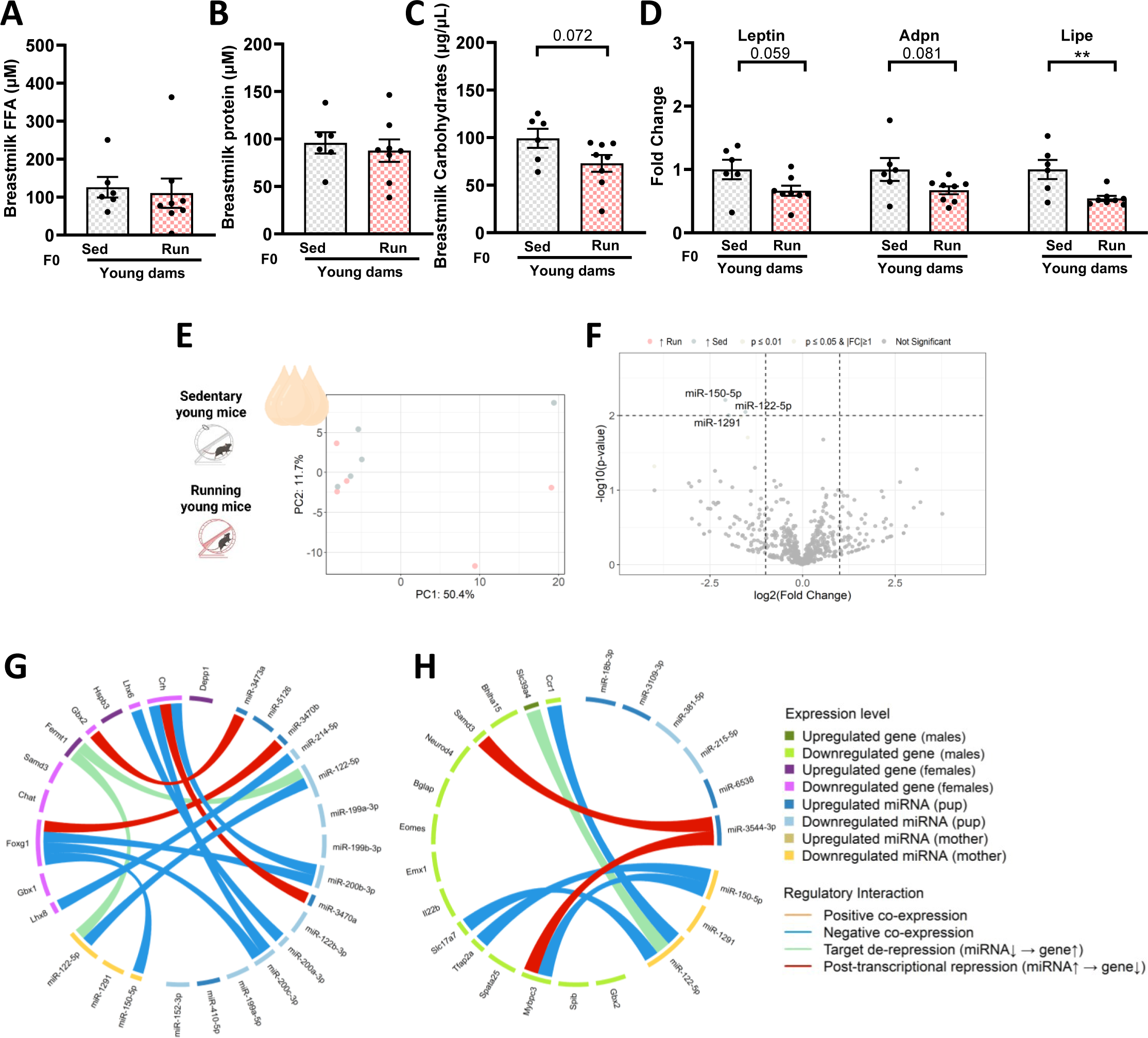
Impact of pre-mating exercise on breastmilk and hypothalamic regulatory network. **A-F**) Analysis of breastmilk content. **A,** Breastmilk free fatty acids concentration. **B,** Breastmilk protein concentration. **C**, Breastmilk carbohydrates concentration. **D**, Gene expression of maternal subcutaneous fat. **E**, PCA of breastfeeding miRNA profile of sedentary and pre-mating dams. **F**, Volcano plots of differentially expressed miRNAs from breast milk of sedentary and pre-mating exercise dams. **G-H**) Regulatory networks. **G**, Regulatory network integrating DE genes in the female offspring hypothalamus with miRNAs from the hypothalamus and maternal breast milk. **H**, Regulatory network integrating DE genes in the male offspring hypothalamus with hypothalamic and maternal breastmilk miRNAs. Unpaired t-test was used in **C**-**F**. Bar graphs show mean ± SEM with individual mice represented as dots. *P≤0.05, **P≤0.01, ***P≤0.001. **A** was created using Biorender.

Beyond its nutritional value of breast milk, breast milk has recently been identified as a rich source of miRNAs that could impact offspring metabolism [45]. Thus, we also analyzed both miRNA content from milk and hypothalamus to investigate whether breast milk miRNA influences the hypothalamus and whether this effect can be modified by maternal pre-mating exercise. Using a threshold of |fold change| > 2 and p < 0.05, we identified 3 downregulated miRNAs (miR-150-5p, miR-122-5p and miR-1291) in breastfeeding milk by pre-mating exercise dams (Fig. 6E-F). MiR-150-5p has been reported previously in the human milk content and associated to immune responses [46] and milk of mothers of preterm compared term infants [47] whereas miR-122-5p and miR-1291 has not been previously described in milk content, but are associated to inflammatory processes [48–50]. To identify how milk miRNAs potentially are coupled to hypothalamic regulation in the offspring, we mapped pairwise comparisons between differentially expressed hypothalamic genes/miRNAs and differentially expressed miRNAs in milk. The results show that these 3 downregulated milk miRNAs may modulate offspring hypothalamic networks involved in neurodevelopment in a sex-dependent manner (Fig. 6G-H).

## DISCUSSION

Here we report that regular pre-mating running impacts offspring phenotype during lactation period. Moreover, we uncovered that the effects of regular pre-mating running on offspring body weight and composition after weaning depend on parental age and offspring sex. Multiple rodent studies have shown the intergenerational effect of physical activity on offspring metabolism. Although the majority of studies underscore the protective effects of both maternal and paternal exercise against metabolic dysfunction in adulthood, some inconsistencies remain [2]. Therefore, we identified that parental age may explain some of the heterogeneity regarding body weight and composition observed in studies on the intergenerational effects of physical activity [2].

Energy homeostasis is conserved in mice [51]. Given our previous observations that energy demand and body composition change throughout mouse male life [31], and considering that metabolic differences between males and females have been described at baseline and in response to exercise [52,53], it was first important to evaluate how energy balance and body composition from young and old dams and sires in response to voluntary exercise. Whereas energy intake was homeostatically regulated when energy demands were high both in young and old males and females, only the young females were resistant to fat loss induced by running. Indeed, the same effect on fat mass regulation was observed in both female and male mice during the second week of running [54]. Interestingly, fat loss linked to caloric restriction was also observed aged female mice and humans, but not in young females [55]. The authors suggested that the activation of relevant pathways by endogenous estrogen may underlie the metabolic resistance of females to calorie restriction [55]. Although it remains unclear whether the same effect occurs in the voluntary wheel-running approach, our findings highlight the importance of investigating not only sex but also age-related adaptations to exercise.

From a biological perspective, many couples are having children at a time when female fertility is declining, with a significant decrease beginning at age 25 and accelerating after age 35 [56]. Ageing not only contributes to a natural increase in fat accumulation in the individual but also in the offspring [24,26]. In line with this, we observed those offspring from older parents weighed more than those of young parents, gained more fat during youth, and exhibited a stronger acute susceptibility to HFD detrimental effects in females, highlighting parental age as a likely contributing factor to the obesity pandemic. In addition to parental age-related weight gain during youth, we observed that parental age interacts with regular pre-mating parental exercise to influence offspring body weight and composition. Considering the null effects of maternal exercise alone, before and during pregnancy, on offspring body composition in mice after weaning, even when voluntary wheel running was performed [57], these results may highlight the context-dependent nature of pre-mating parental exercise on offspring body weight and composition after weaning. However, given that expression changes in the subcutaneous fat and hypothalamus in male offspring in response to regular pre-mating parental exercise, and the beneficial effects of maternal exercise on offspring metabolism frequently emerge during adulthood [45], further longitudinal studies are warranted to investigate the effect of parental age and pre-mating parental exercise on offspring adulthood.

During lactation, we did not observe differences in offspring body weight at 14 and 21 days of age, as observed previously in offspring rats from exercised dams prior to and during mating, pregnancy, and nursing [58]. In contrast, we observed that pre-mating exercise accelerated the body weight changes before weaning, which may underlie a role of regular pre-mating parental exercise during lactation. Indeed, offspring from pre-mating exercised parents displayed less subcutaneous fat and visceral fat than those of sedentary parents at 9 days old. Moreover, the tibia length was slightly smaller in those offspring from exercised parents, being more pronounced in females, suggesting a likely role of pre-mating parental exercise on offspring development. Interestingly, it has been previously speculated that parental exercise may have resulted in developmental delays, given the delayed eye opening in offspring of dams exposed to exercise for 2 weeks before mating, 18 days during breeding, and 10 days postpartum, with sires also allowed to use running wheels during 18 mating days [32]. These changes during lactation are independent of maternal care since pre-mating exercise in dams had no effect on nest-building behavior or the time spent with pups, both of which are used as indirect indicators of maternal care. Whether parental age impacts maternal care remains to be investigated.

Since maternal exercise during lactation may improve breast milk composition and metabolome potentially impacting on offspring adiposity [45,59] and breastmilk miRNAs content has been associated with both maternal and offspring body composition [60,61], we next evaluated whether pre-mating maternal exercise could induce long-lasting effect of subcutaneous fat which, in turn, would affect differentially in the breastfeeding milk content offspring adiposity. Pre-mating maternal exercise modestly influenced gene expression in subcutaneous fat and produced minor changes in milk nutritional content and miRNA levels at 9 days postpartum. In line with this, it has been reported that maternal exercise during lactation induced only minor alterations in the metabolism of mammary epithelial cells in lean mice. However, it partially reversed the obesity-induced changes in mammary gland metabolism and milk fatty acid composition [62]. Our premise is that the specific timing of maternal exercise may differentially affect mammary metabolism in older dams and offspring phenotypes, which will be addressed in future studies. Evidence suggests that small extracellular vesicles and their miRNA cargos of breastfeeding milk may resist offspring gastrointestinal passage and cross the offspring blood-brain barrier [63–66]. Surprisingly, the three downregulated milk miRNAs may modulate offspring hypothalamic networks involved in neurodevelopment in a sex-dependent manner, which may open new avenues for understanding how parental exercise might influence offspring brain development.

Several limitations of this current study should be considered. Although wheel running approach constrains the ability to regulate volume, intensity and timing of running, it mimics interval training and active lifestyle in humans [67] and induces similar changes to treadmill approach regarding body composition changes without introducing a stress response [68,69]. In mice, the energy balance is primarily regulated by total energy expenditure, specially at ∼22°C [70,71], which differs from humans, who live within their thermoneutral comfort zone [70,72]. Since our experiments were not conducted at thermoneutrality, the translatability of these preclinical findings is questionable. It is important to note that this study is not powered, and the offspring sample size was limited because the dams were considered the statistical units [33]. Moreover, the estrous cycle of both female dams was not monitored during testing, and this may have impacted on the outcomes during running period and pregnancy [73,74].

In conclusion, our work underscores the need to consider parental age when evaluating offspring phenotypes, highlights that regular parental exercise even before mating can influence offspring adiposity during the period of exclusively breastfeeding milk period and uncovers the importance of studying both female and male offspring to show the sex specific effects in offspring.

## AUTHOR CONTRIBUTIONS

Conceptualization, P.S.; Methodology, P.S; Formal analysis, C.M, C.C, J.V and P.S; Investigation, C.M, J.V, and P.S.; Visualization, C.M and P.S., Writing–original draft, C.M, J.V and P.S; Writing – review & editing, C.M, C.C, J.V, and P.S; Supervision, P.S; Funding acquisition, P.S.

## ACKNOWLEDGEMENTS

Anne Jørgensen, Lene Foged and Ida Holm are acknowledged for their technical assistance. This study was supported by a research grant from Lundbeck Foundation (grant ID R380-2021-1300) award by P.S, the Novo Nordisk Foundation (grant ID 0059436) and the Centre for Physical Activity Research (CFAS), which is supported by TrygFonden (grants ID 101390, ID 20045, ID 125132, and ID 177225).

## LEAD CONTACT

Further information and requests for resources and reagents should be directed to and will be fulfilled by the Lead Contact, Paula Sanchis.

## DECLARTION OF INTERESTS

The other authors declare no competing interests. All authors gave their approval for the current version to be published.

## RESOURCE AVAILABILITY

### Materials Availability

This study did not generate new unique materials.

### Data and Code Availability

The transcriptomic raw RNA-seq data have been deposited at the Sequence Read Archive Database under BioProject accession PRJNA1301823

## SUPPLEMENTAL INFORMATION TITLES AND LEGENDS

Document S1. Supplementary Figures

## METHODS

### Mouse experiments

Mouse experiments complied with the European directives regarding the care and use of experimental animals, approved by the Danish Animal Experiments Inspectorate (2020-15-0201-00599) and were conducted in adherence with ARRIVE guidelines. Two cohorts of mice were purchased from Janvier Labs and maintained at the animal facility of the University of Copenhagen, Denmark. In both cohorts, 7-week-old C57BL/6NRj mice (young parents), 20 females and 10 males, and 26-week-old C57BL/6JRj mice (old parents), 20 females and 10 males, were scanned using EchoMRI Body Composition Analyzer to determine the body composition and assign mice equally, according to body fat, into an active (with a functional 4¾” x 4.5” running wheel from Starr, Life Sciences Corp.) or a sedentary (with a blocked running wheel) group. All animals used in this study were single-housed at 22±2C with a 12-h light/dark circle, had *ad libitum* access to water and were provided with nesting, bedding material, small plastic house and cardboard tube. Mice were regularly feed *ad libitum* chow diet (SAFE D30, Safe Diets, 3.389 Kcal/gram; 22% protein, 5.3% fat, and 50.8% carbohydrates; www.safe-lab.com).

#### Cohort 1

Food intake, running distance and body weight were measured daily and the body composition weekly during running period at ZT10-12. Energy expenditure and energy balance was estimated using virtual calorimeter v1.82 [75]. After 5 weeks of voluntary wheel running, mice from each group were mated separately by putting together 2 females and 1 male per cage and the litter number and size of 4 groups were recorded. Males were removed from cages after 19 days of mating. In case that both females from same cage gave birth the same day, the mean of total and alive pups per female was calculated. Siblings from a given litter remained together and offspring body weight was monitored from 2-week-old and offspring body composition was measured weekly after weaning. To minimize the littermate effect on the results, we analyzed body weight and composition after weaning from one male and one female from litters with more than six pups selecting the mice based on the mean body weight on day 14, separated by sex. Next, female offspring were fed *ad libitum* HFD (D12492, 5.24 kcal; 20% energy from protein, 60% energy from fat and 20% energy from carbohydrates, www.researchdiets.com) for 14 days. Male offspring were subjected to voluntary wheel running to evaluate the intergenerational effect of running on offspring performance and phenotype. To conduct this experiment, additional male per litter was added based on food intake and body weight on day 0 of the experiment.

#### Cohort 2

Running distance was monitored throughout the experiment whereas body weight and body composition were assessed in all mice at baseline (one week after arrival) and after 6 weeks of running (day 47). Given that behavioral studies were sequentially conducted on the 60 mice from day 15 to day 26, the running period was extended to 6 weeks instead of 5 to minimize potential behavioral testing effects on mating. A polygamous trio was selected as the breeding strategy, as mentioned above, for 5 days to avoid birth timing discrepancies. Each female was then isolated in a separate cage to accurately record which pups belonged to which female. During the subsequent 3-week gestation period, birth dates and the number of pups in each litter were recorded. On postnatal day P2, litters were recounted to assess number of pups lost, potentially as a result of maternal cannibalistic behavior, and on the same day, a random selection of pups was euthanized to standardize all litters to six pups [33]. On postnatal day P7, maternal behavior was analysed by recording the cages for one hour from 3-5 PM. To acclimate the animals to the recording environment, the cages were removed from the racks and placed in the recording area 24 hours prior to filming. On postnatal day P9, both the offspring and the dams were euthanized, and maternal breastmilk and maternal and pup tissues were collected.

From voluntary wheel running protocol, specific datapoints were excluded both for sedentary and running mice owing to measurement errors, then mean from day before and after were used. In case of daily measurements could not be taken, last and first day of missing data was averaged. Several datapoints were also missed owing to technical problems with the counters. Statistical analyses were performed in IBM SPSS Statistics v29.0.1.0 and statistical analysis information for each graph can be found in the Figure legends. No statistical methods were applied to predetermine the sample size for experiments. C.M. and P.S. were aware of the group allocations in different experiments.

### Behavioral tests

The parental behavioral testing was conducted in the behavioral room at ZT9-12, placing the mice to the different apparatuses. Mice were habituated to the behavioral room one week in advance and behavioral battery started with y-maze, followed by elevated plus maze and ended with open field, with several days of rest between tests for each animal.

Y-maze test assesses short-term spatial working memory [76]. Mice were placed at the end of one arm of a Y-shaped maze and allowed to explore freely for 8 min. The number of arm entries and alternations were recorded, with an alternation defined as successive entries into each of the three arms. The arms entries were noted in situ during the test and number of correct alternations was calculated using a custom Python script.

The Elevated Plus Maze (EPM) evaluates anxiety-like behavior in a novel and unrestricted environment [77] by analyzing the number of closed- and open-arm entries and the time in each arm for 5 min in Ethovision XT version 11.5 (Noldus) software. In cases where recording issues occurred, the respective mouse was excluded from the analysis.

The open field (OF) test was used to assess general locomotor activity and anxiety-like behavior. Mice were placed in a large square arena (50x50 cm) with clear walls and allowed to explore for 5 min. Total distance travelled, time spent in the center and time spent in the periphery was recorded and analysed using the Ethovision XT version 11.5 (Noldus) software. In cases where recording issues occurred, the respective mouse was excluded from the analysis.

Nesting behavior reflects general health and parental ability to care for and keep pups warm [38,39]. It was conducted overnight placing a 3-gram cotton fiber material (Ancare Corporation NES3600) at ZT11-12 into home cages of female mice during to evaluate the nest and weigh the untorn residue in the next day at ZT4 using a 5-point nest-rating scale [38].

To evaluate integrated anxiety z-scores, we applied behavioral z-score normalization to open field and elevated plus maze data in males, and to nesting behavior in females, as described by Bucknor and collaborators [36].

Maternal presence in the nest was assessed on postnatal day 7 over a 1-hour observation period. Dams were recorded as present in the nest at 3-minute intervals. To minimize disturbance, observations were conducted in the animals’ home cages, which were open-top to allow recording. If a dam was found outside the cage during the observation period, the experimenter briefly entered the room to gently return her to the cage. These intervention periods were excluded from the analysis. Dams that spent more than 50% of the observation period outside the cage were excluded from the analysis.

### Breast milk collection and nutritional content

Milk was collected from 2^nd^ cohort dams at P9. Briefly, dams were separated from their pups for ∼3 hours in a transparent-housed closed to pup housed to maintain the sensory stimulus but avoiding breastfeeding milk from ZT6. At ZT9, mums were intraperitoneally injected with 2 IU/kg of oxytocin (Orifarm 10 IE) and rapidly anesthetized by inhalation with 3.0–3.5% isoflurane in oxygen. Breast milk was carefully sucked into the syringe tube by a vacuum using a home-made device, as described by Klein and collaborators [78], and stored at -80C. One aliquot was using for evaluating the miRNA content and the other for analyzing the nutritional content For the nutritional content, the aliquot was divided into three aliquots for the analysis of carbohydrates, free fatty acids (FFA), and proteins. Carbohydrate content was analysed using three times 1:50 dilutions prepared according to the Sigma-Aldrich Total Carbohydrate Assay kit protocol (Cat number MAK559). For FFA analysis, 1:50 dilutions were prepared in 1.5 mL Eppendorf tubes containing 176 μL of the kit’s assay buffer, 10 μL of 5% Triton X-100, 10 μL of 5% isopropanol, and 4 μL of milk. Tissue was homogenized with plastic pestles, and the analysis was performed using the Sigma-Aldrich Free Fatty Acid Assay Kit (Cat number MAK466). Protein concentration of the individual samples was determined by using the PierceTM BCA protein assay kit relative to a Bovine Serum Albumin (BSA) standard. Breastmilk samples were diluted 1:100 in distilled water, following the Thermo Scientific PierceTM BCA Protein Assay Kit protocol (Cat number 23225). After incubation, the absorbance for both the standards and the samples was determined using a TECAN Sunrise™ reader. All analyses were carried out using the SUNRISE ELISA reader, and results were processed and analysed in Excel.

### Tissue harvesting

Pups from the 1^st^ cohort were sacrificed by decapitation after the running period (males) or after HFD feeding (females). The hypothalamus was isolated, snap-frozen immediately in liquid nitrogen, and stored at −80 °C. The right hind limbs were removed and stored in 5 mL tubes filled with tap water for one week to allow skin removal. Tibia length was then measured using a manual vernier caliper.

Dams from the 2^nd^ cohort were sacrificed by decapitation following breast milk collection, and samples of subcutaneous fat were collected. Pups from 2^nd^ cohort were weighed and euthanized alternately from different litters born on postnatal day P9 to ensure consistent endpoint timing across animals. Blood was collected from the trunk, and peripheral tissues, including subcutaneous fat, visceral fat, and brown adipose tissue from the neck were weighed, snap-frozen in liquid nitrogen, and stored at −80 °C. The hypothalamus was also snap-frozen immediately for later analysis. The right hind limbs were removed, and the same procedure described above was followed.

### Pups sex determination

To determine the sex of the offspring from 2^nd^ cohort, DNA was extracted and subsequently analysed using quantitative PCR (qPCR) targeting the *SRY* gene, which is exclusive to the Y chromosome. The presence of *SRY* expression indicates male offspring, while its absence indicates female offspring. DNA extraction was performed following the manufacturer’s instructions provided with the Omega Biotek E.Z.N.A.® Circulatory Tissue DNA Kit (Catalog No. D3396-02, 200 preps), which was used for efficient DNA isolation from tissue samples. qPCR was then conducted to amplify the *SRY* gene, allowing for sex determination of the offspring.

### Gene expression analysis

Dam subcutaneous fat from 2^nd^ cohort and offspring subcutaneous, brown and hypothalamus from 2^nd^ cohort were used for RNA extraction. RNA was extracted using TRIzol® reagent (Invitrogen, Cat# 15596018). Tissue samples were homogenized in 1 ml of TRIzol® using a Tissuelyser (Qiagen) with a metal bead at 25 Hz for 2 cycles of 1,5 minute. After homogenization, the samples were left to settle for 5 minutes at room temperature, then centrifuged at 12000 g for 10 minutes at 4°C. The TRIzol-containing supernatant was carefully transferred to a new Eppendorf tube, ensuring that neither the pellet nor the upper transparent fat layer was disturbed. To this new tube, 200 μl of chloroform (Sigma Aldrich, Cat# 34854) was added, and the samples were shaken for 20 seconds and incubated for 5 minutes at room temperature. Samples were then centrifuged at 12000 g for 10 minutes at 4°C to separate the phases. The upper aqueous phase, containing RNA, was carefully collected and mixed with an equal volume of ice-cold isopropanol (Sigma Aldrich, Cat# 34863). The mixture was vortexed, incubated at RT for 10 minutes, and then centrifuged at 12000 g for 15 minutes at 4°C to pellet the RNA. The RNA pellet was washed twice with ice-cold 75% ethanol (Sigma Aldrich, Cat# 64-17-5), and the samples were centrifuged at 8000 g for 5 minutes at 4°C. After drying, the RNA was resuspended in nuclease-free water (Invitrogen, Cat# 10977-035) and stored at -80°C. RNA concentration was measured using a Nanodrop spectrophotometer and reversed transcribed using cDNA Reverse Transcription Kit (1 μl Reverse Transcriptase (RT), 2μl 10x RT buffer, 0,8 μl dNTP, 2μl 10x RT random primers, 1μl RNase inhibitor and 3,2μl ultrapure nuclease free water, Cat# 4368813, Thermo Fisher Scientific) following the instructions from the manufacturer and run at the following program for reversed transcription (25 °C for 10 min., 37 °C for 120 min., 85 °C for 5 min, 4 °C ∞).

Quantitative PCR (qPCR) were performed with PowerUp^TM^ SYBR^TM^ Green Master Mix (Cat# A25780, ThermoFisher Scientific), 300 nM of forward and reverse primers and using ViiA 7 Real Time PCR system (Applied Biosystems) for amplification following parameters: hold stage at 95 ℃ for 20 secs, followed by 40 PCR cycles consisting of denaturation at 95 ℃ and annealing, extension and fluorescence reading at 60 ℃. Relative quantification using ΔCt with a fold change normalization at the sedentary group was calculated. *Gadph* was used as housekeeping gene for all tissues. Most sequences were obtained from PrimerBank[79]. Primers were synthetized by TAG Copenhagen (see below). Statistical analyses were performed in IBM SPSS Statistics v29.0.1.0.

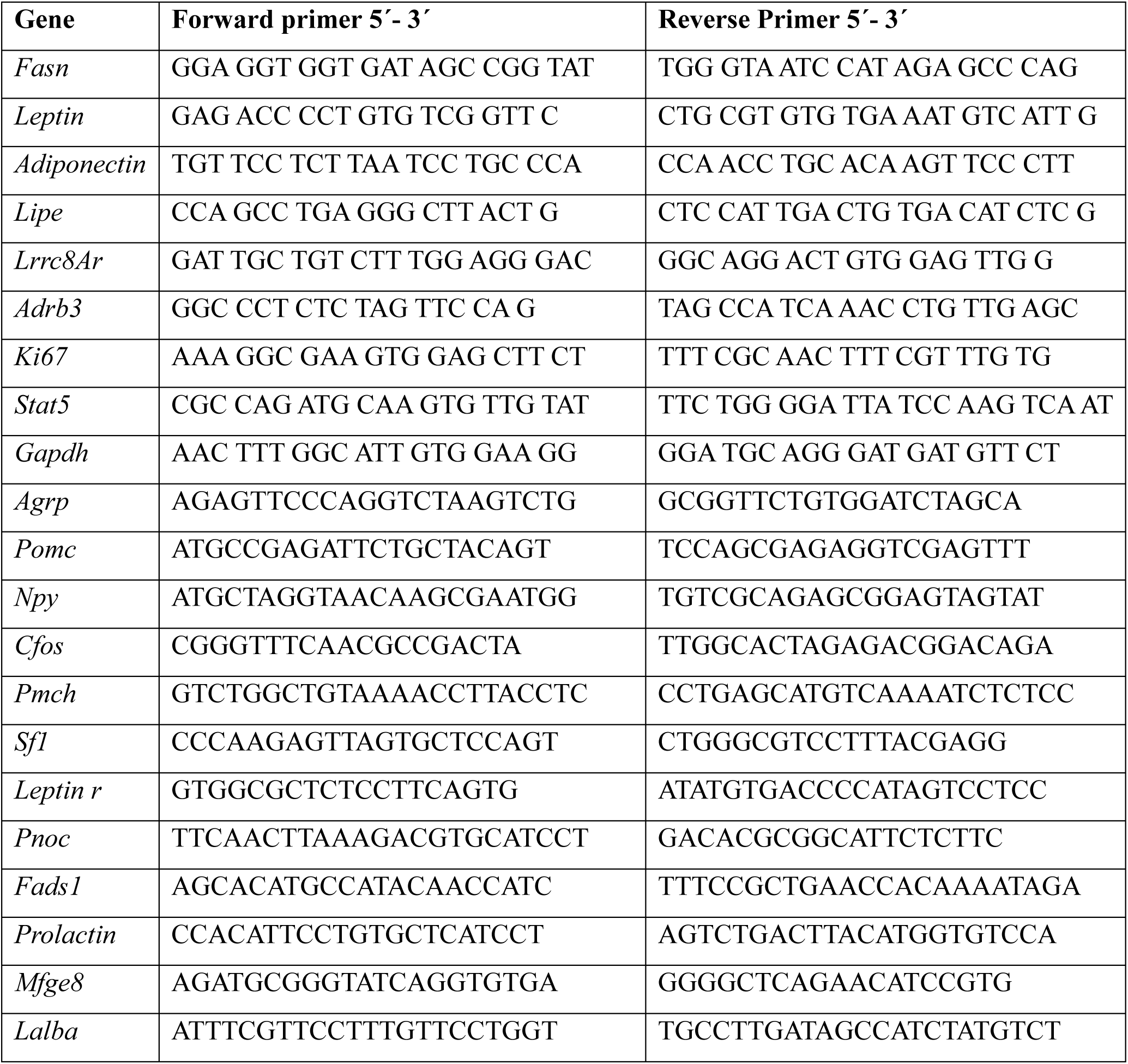

### Bulk transcriptomic profile by RNA sequencing

We selected litters from 2^nd^ cohort with an equal number of male and female offspring. Within each litter, individuals whose weight at P9 was closest to the mean were chosen. RNA from offspring hypothalamus were obtained using Trizol-phenol method as previously described and sent to Genewiz from Azenta Life Sciences for sequencing. Stranded RNA-seq libraries were prepared for all high-quality RNA samples using mRNA with poly A tail enrichment. The libraries were then sequenced in Illumina, PE 2x150bp, generating ∼50 million paired-end reads per sample. Quality control of the raw sequencing data was performed with SOAPnuke v2.1.5 with default options [80]. The reads for each of the samples were mapped to the *Mus musculus* genome assembly GRCm39 using STAR v2.7.11b with default parameters [81]. The gene counts were retrieved using featureCounts v2.0.6 [82], and then built into a matrix. Low-count genes were filtered based on the count-per-million and sample library size (the gene is expressed in at least one replicate). The expression abundance was quantified using cufflinks v2.2.1 [83], retrieving the fragments per kilobase of transcript per million mapped reads (FPKM) values.

### Transcriptomic deconvolution

Bulk RNA-seq data from hypothalamic samples were variance-stabilized using DESeq2 v1.46.0 [84], and estimated cell-type deconvolution was performed with MuSiC algorithm [85]. As reference, we used the HypoMap single-cell RNA-seq atlas [86], providing curated annotations of hypothalamic cell populations. MuSiC [85] was applied independently to female and male datasets to estimate cell-type proportions based on the single-cell reference. The resulting weighted proportion matrices were merged, normalized per sample, and averaged within each experimental group. Data processing and visualization were carried out in R (v4.3.2) using the tidyverse R library [87], generating a stacked bar plot to summarize hypothalamic cell-type composition under each condition.

### Dam breast milk and hypothalamic miRNA content

Dam breast milk and a hypothalamic tissue from male and female littermates were used for obtaining copurification of miRNA content and total RNA using miRneasy Qiagen kit (Cat number #1071023). Samples were sent to Genewiz from Azenta Life Sciences for sequencing. Small RNA-seq libraries were prepared for all high-quality copurification of miRNA content and total RNA samples using Small RNA-seq. The libraries were then sequenced in Illumina, PE 2x150bp. An estimated data output of ∼20 million paired-end reads per sample were generated.

Following the workflow described by Xu and colleagues [88], raw read quality was assessed with FastQC v0.11.9 [89], adapters were trimmed using Cutadapt v3.4 [90], and filtered reads were aligned to mature *Mus musculus* sequences from miRBase v22 [91] using Bowtie v1.3.1 [92]. Alignment files were processed with SAMtools v1.17 [93], and counts were aggregated into two matrices, one for pups and one for milk. Differential expression analysis for both RNA-seq and miRNA-seq was performed in R v4.3.2 using DESeq2 v1.46.0 [84]. Variance-stabilizing transformation (VST) was applied for visualization and exploratory analysis, including PCA, volcano plots, and heatmaps. Functional enrichment of DEGs and predicted targets of differentially expressed miRNAs retrieved with multiMiR [94]was conducted with clusterProfiler [95] using Gene Ontology Biological Processes and *Mus musculus* annotations from org.Mm.eg.db [96]. Significance was evaluated with Benjamini–Hochberg adjusted *p* ≤ 0.05, and fold enrichment was calculated as the ratio of observed to expected genes per term.

### Hypothalamic Regulatory Networks

A custom circular network diagram was developed to visualize regulatory interactions between RNAs and miRNAs from dams and their offspring. Male and female offspring were conducted separately. Raw count matrices for mRNA and miRNA (both from breastmilk and pup hypothalamus) datasets were imported and pre-processed separately for each experimental group. Both mRNA and miRNA samples were normalized and processed using DESeq2 v1.46.0 [84] with identical design formulas and statistical thresholds applied to all datasets (*p value* ≤ 0.01, |log₂FC| ≥ 1). Differentially expressed transcripts were subsequently classified as upregulated or downregulated.

To explore post-transcriptional regulatory relationships, predicted and experimentally validated miRNA–mRNA interactions were retrieved using the multiMiR database (v2.1) [94] for *Mus musculus*. Interactions were retained only when both the miRNA and its target gene were significantly regulated within the respective datasets. Shared miRNAs between dams and pups were cross-compared to identify concordant or opposite regulatory trends. A global regulatory network was then assembled, integrating maternal miRNAs, pup miRNAs, and mRNAs. The final network was visualized as a chord diagram using the *circlize* package [97]with nodes color-coded by expression direction and chords representing distinct interaction types (co-expression, de-repression, or post-transcriptional repression).

## SUPPLEMENTARY FIGURE LEGEND

**Suppl. Figure 1. Running increases energy expenditure but does not affect overall energy balance. A-C**) Regular exercise in young and old males. **A**, Schematic. **B**, Energy expenditure. **C**, Energy balance. **D-E**) Regular exercise in young and old females. **D**, Schematic. **E**, Energy expenditure. **F**, Energy balance. **G**-**L**) Changes in body weight and body composition during pregnancy of sedentary and pre-mating dams. **G**. Body weight of pregnant females. **H**, Body weight changes of pregnant females. **I**, Body weight at weaning of pregnant dams with surviving pups. **J**, Fat mass of pregnant dams with surviving pups. **K**, Lean mass of pregnant dams with surviving pups. **L**, Body composition changes of pregnant dams with surviving pups. Two-way repeated-measures ANOVA was used to assess running (*), age (#) and its interaction ($) in **B**-**C** and **E**-**F**, but if sphericity and/or normality assumptions were significant, then repeated-measures GEE was conducted. Two-way ANOVA was used in **G**-**L**, but if normality test and/or homogeneity of variances were violated, GzLM comparisons was used. Time-course data are presented as mean ± SEM, and bar graphs show mean ± SEM with individual mice represented as dots. *P≤0.05, **P≤0.01, ***P≤0.001. **A** and **D** were created using Biorender.

**Suppl. Figure 2. Further phenotypic and molecular analysis related to Fig. 3. A, Schematic. B-D**) Gene expression changes in female offspring tissues. **B**, Subcutaneous Fat. **C**, Visceral Fat. **D**, Hypothalamus. **E**-**L**) Phenotypic analysis of offspring male in response to parental pre-mating exercise and parental age. **E**, Fat mass, **F**, Lean mass. **G**, Body weight. **H**, Body weight changes. **I**, Estimated energy expenditure. **J**, Estimated energy balance. **K**, Tibia length. **L**, Tissues. **M**-**N**) Gene expression changes in male offspring tissues. **M**, Visceral Fat. **N**, Hypothalamus. Two-way ANOVA was used to assess parental running, parental age (#) and its interaction in **B**-**D** and **L-N** and three-way ANOVA was used for evaluating the effects of pre-mating parental exercise, parental age, offspring running (*), and its interaction in **H**-**K**. However, GzLM was used if normality and/or homogeneity were violated. Three-way repeated-measures ANOVA was used to assess offspring running, parental age, pre-mating parental exercise and its interaction ($) in **E**-**G**, but if Mauchly’s sphericity test and/or Shapiro-Wilk’s Normality test was significant, then repeated-measures GEE was conducted. Bar graphs show mean ± SEM with individual mice represented as dots. *P≤0.05, **P≤0.01, ***P≤0.001. **A** was created using Biorender.

**Suppl. Figure 3. Second cohort of parents related to Fig. 4. A-E**) Regular exercise period. **A**, Schematic. **B**, Distance. **C**, Fat mass changes. **D**, Lean mass changes. **E**, Body weight changes. **F-I**) Pregnancy outcomes in response to parental age and pre-mating parental exercise in 2^nd^ cohort. **F**, Table of pregnant females. **G**, Pups per littermate. **H**, Survived pups. **I**, Females pup rate. Two-way ANOVA was used for evaluating the effects of parental running (*), parental age (#) and its interaction ($) in **B** whereas **t**hree-way ANOVA was used for evaluating the effects of parental running, parental age, parental sex (^) and its interaction ($ for parental running*parental age, ¤ for parental running* parental sex, • for parental age* parental sex, and & for triple interaction) in **C**-**E**. However, GzLM was used if normality and/or homogeneity were violated. Unpaired t-test was used in **I** and no statistical analyses were applied in **G**-**H** given only 1 old females was pregnant. Bar graphs show mean ± SEM with individual mice represented as dots. *P≤0.05, **P≤0.01, ***P≤0.001. **A** was created using Biorender.

**Suppl. Figure 4. Further molecular and bioinformatic analysis related to Fig. 5. A-C**) Offspring gene expression changes in response to pre-mating parental exercise. **A**, Schematic. **B**, Brown fat analysis in male offspring. **C**, Subcutaneous fat analysis in male offspring. **A** was created using Biorender. **D**) Distribution of cell proportions estimated from RNA-seq deconvolution. **E-F**) GO biological process enrichments. **E**, GO of DE genes from female offspring. **F**, GO of DE genes from male offspring. Two-way ANOVA was used for evaluating the effects of parental running (*), offspring sex (^) and its interaction in **B**-**C**. GzLM was used if normality and/or homogeneity were violated. Bar graphs show mean ± SEM with individual mice represented as dots. *P≤0.05, **P≤0.01, ***P≤0.001.

**Suppl. Figure 5. Gene expression analysis of maternal subcutaneous fat. A)** The subcutaneous fat of sedentary and running dams was analyzed at 9 days post-partum. T-test was used for assessing the effects of running (*) in all graphs. Bar graphs show mean ± SEM with individual mice represented as dots. *P≤0.05, **P≤0.01, ***P≤0.001.

