## Supplementary material for "Regular parental exercise before mating influences offspring lower adiposity associated to hypothalamic neurodevelopmental changes": Suppl figures

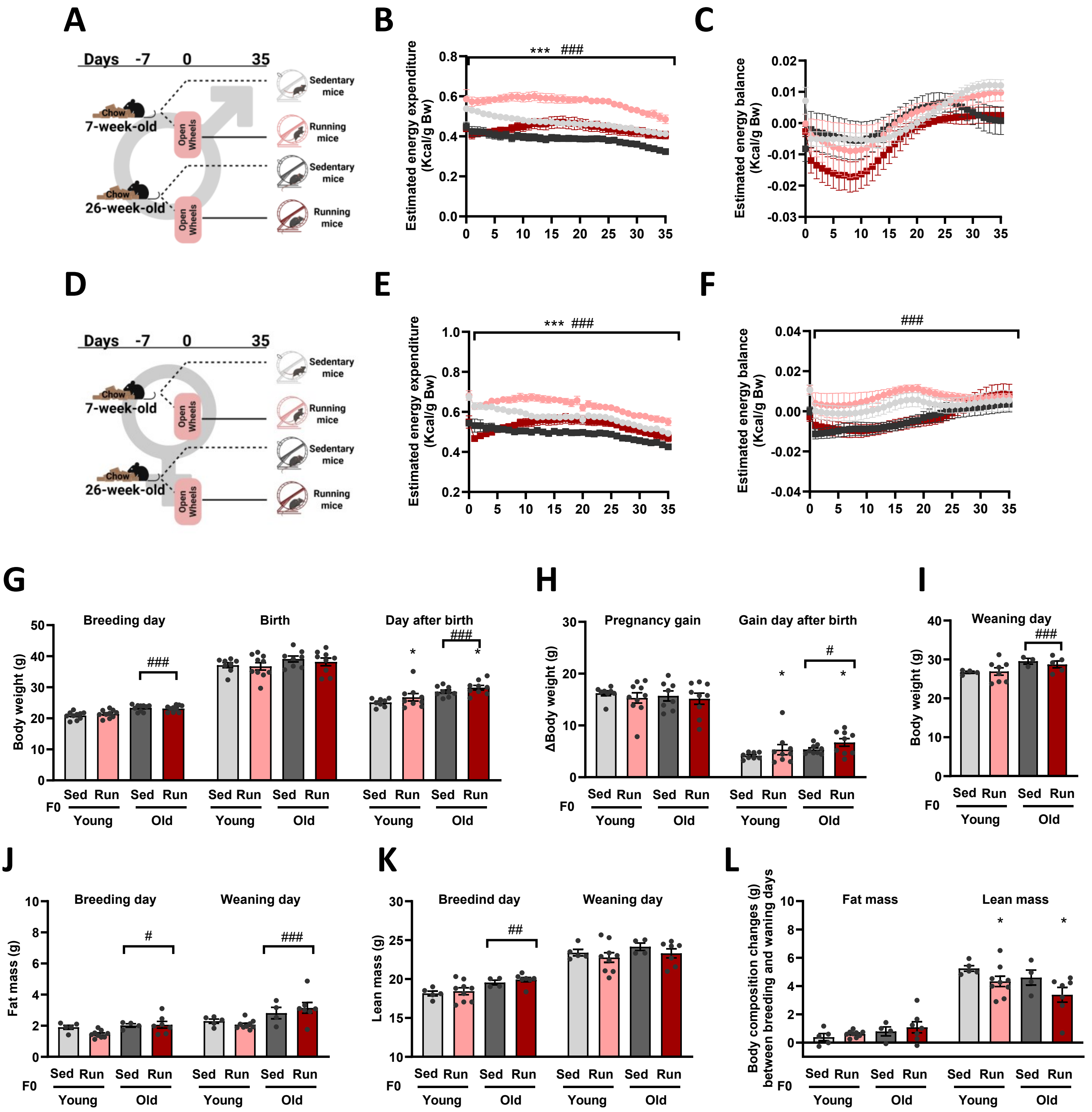
A
Days -7 0 35
Chow
7-week-old
26-week-old
Open Wheels
Sedentary mice
Running mice
B
Estimated energy expenditure (Kcal/g Bw)
*** ###
C
Estimated energy balance (Kcal/g Bw)
D
E
F
###
G
Breeding day
Birth
Day after birth
Body weight (g)
Sed Run Sed Run
F0
Young Old
H
Pregnancy gain
Gain day after birth
ΔBody weight (g)
I
Weaning day
J
Fat mass (g)
K
Breedind day
Weaning day
Lean mass (g)
L
Body composition changes (g) between breeding and waning days
Fat mass
Lean mass

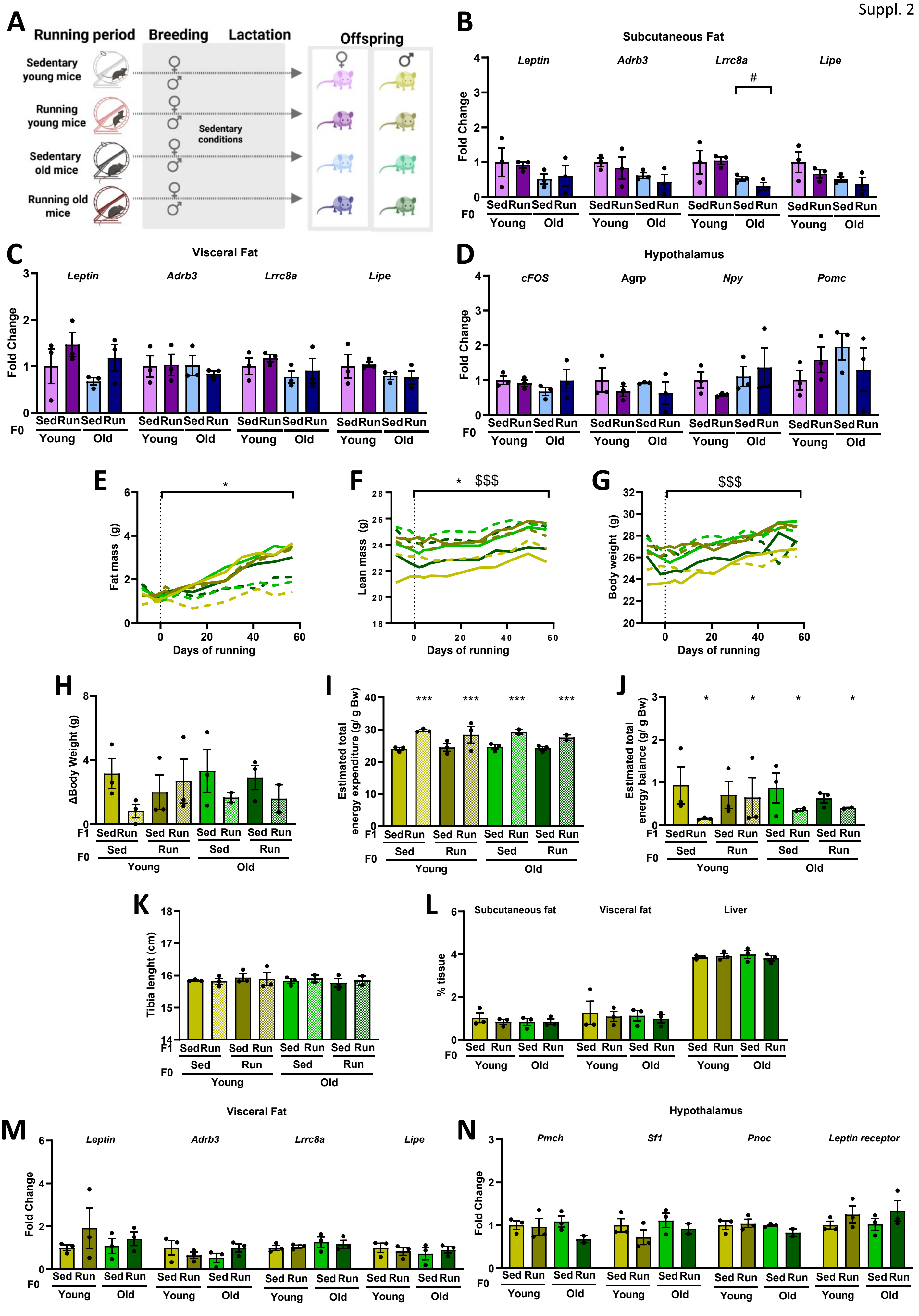
A
Running period
Breeding
Lactation
Offspring
Sedentary young mice
Running young mice
Sedentary old mice
Running old mice
Sedentary conditions
B
Subcutaneous Fat
Leptin
Adrb3
Lrrc8a
Lipe
Fold Change
Sed Run Sed Run
F0
Young Old
C
Visceral Fat
D
Hypothalamus
cFOS
Agrp
Npy
Pomc
E
Fat mass (g)
Days of running
F
Lean mass (g)
G
Body weight (g)
H
ΔBody Weight (g)
F1
I
Estimated total energy expenditure (g/ g Bw)
J
Estimated total energy balance (g/ g Bw)
K
Tibia lenght (cm)
L
Subcutaneous fat
Visceral fat
Liver
% tissue
M
Visceral Fat
N
Hypothalamus
Pmch
Sf1
Pnoc
Leptin receptor

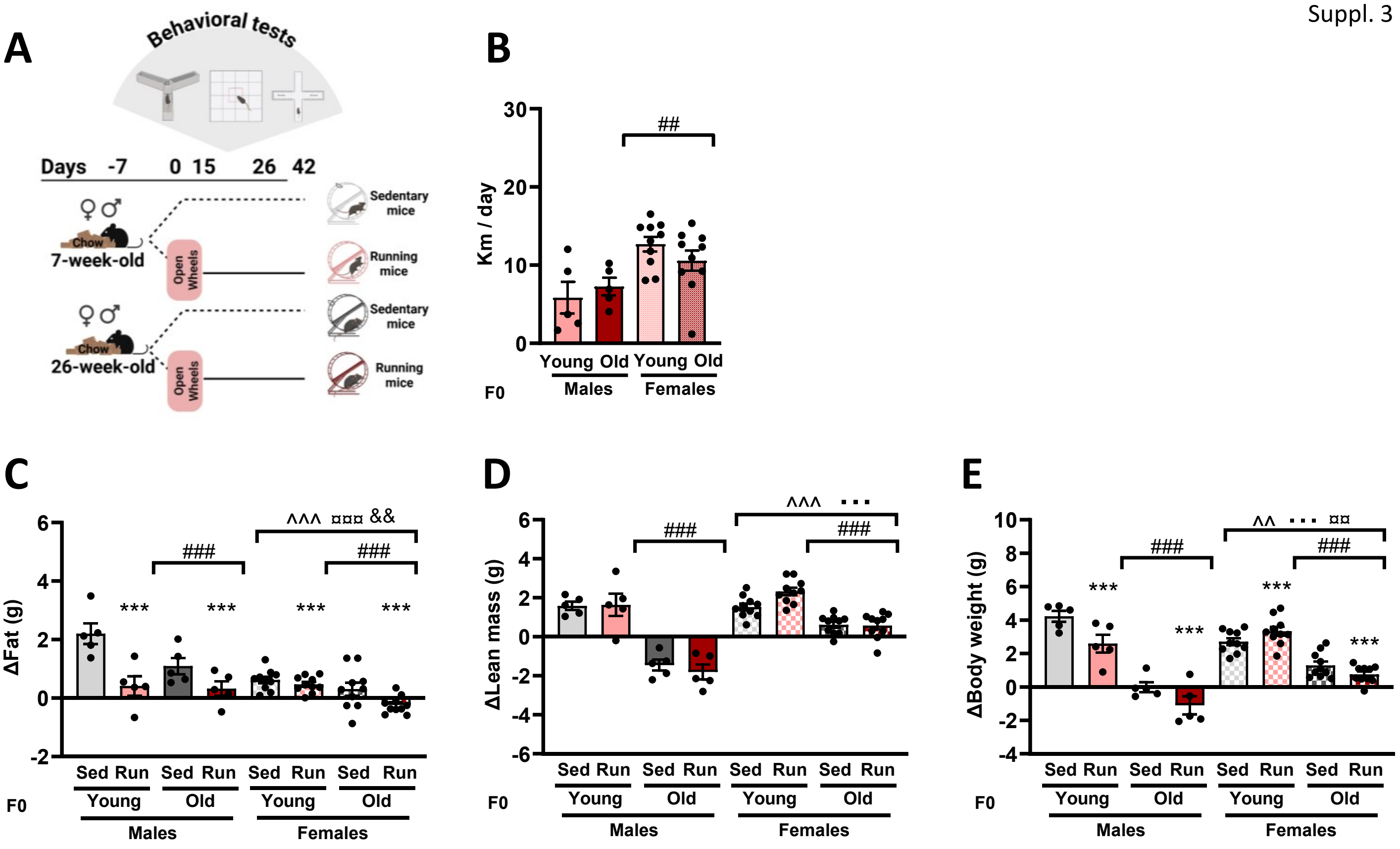


F

| | Total females (n) | Total pregnant (n) |
|---|---|---|
| Young sedentary females | 10 | 7 |
| Young running females | 10 | 10 |
| Old sedentary females | 10 | 1 |
| Old running females | 10 | 4 |

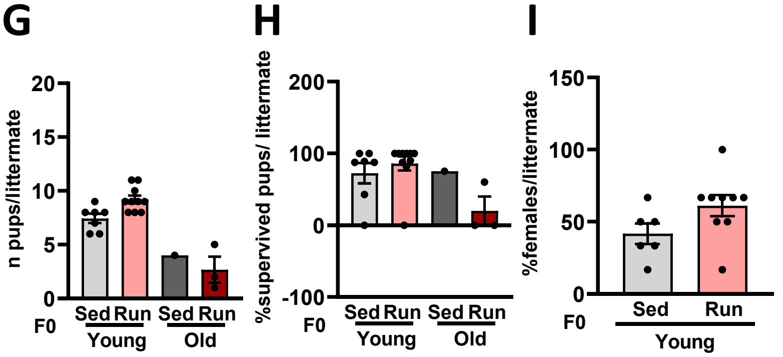

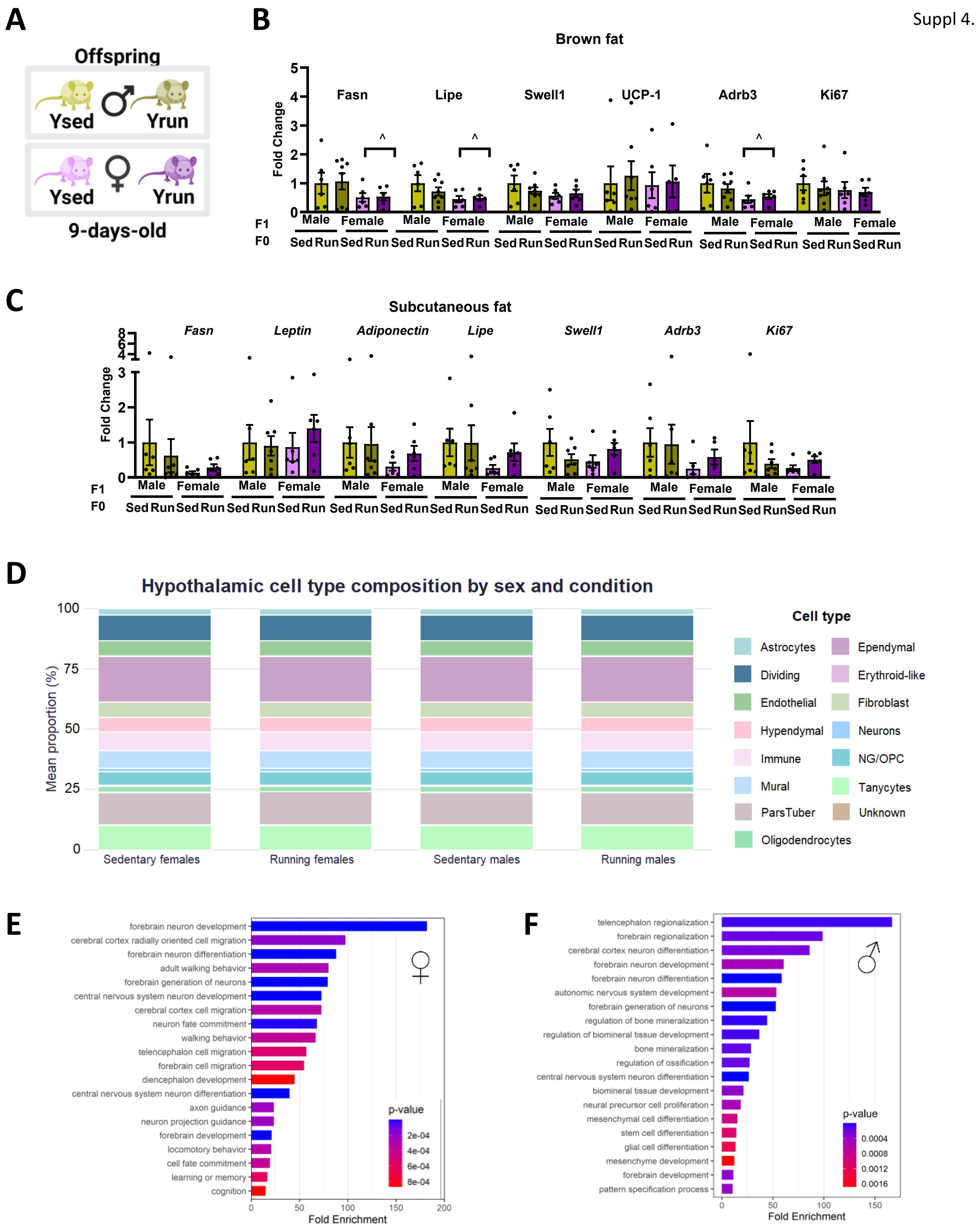
A
Offspring
Ysed
Yrun
Ysed
Yrun
9-days-old
B
Brown fat
Fold Change
Fasn
Lipe
Swell1
UCP-1
Adrb3
Ki67
Male Female
F1
F0
Sed Run Sed Run
C
Subcutaneous fat
Fold Change
Fasn
Leptin
Adiponectin
Lipe
Swell1
Adrb3
Ki67
D
Hypothalamic cell type composition by sex and condition
Mean proportion (%)
Sedentary females
Running females
Sedentary males
Running males
Cell type
Astrocytes
Ependymal
Dividing
Erythroid-like
Endothelial
Fibroblast
Hypendymal
Neurons
Immune
NG/OPC
Mural
Tanycytes
ParsTuber
Unknown
Oligodendrocytes
E
forebrain neuron development
cerebral cortex radially oriented cell migration
forebrain neuron differentiation
adult walking behavior
forebrain generation of neurons
central nervous system neuron development
cerebral cortex cell migration
neuron fate commitment
walking behavior
telencephalon cell migration
forebrain cell migration
diencephalon development
central nervous system neuron differentiation
axon guidance
neuron projection guidance
forebrain development
locomotory behavior
cell fate commitment
learning or memory
cognition
p-value
Fold Enrichment
F
telencephalon regionalization
forebrain regionalization
cerebral cortex neuron differentiation
forebrain neuron development
forebrain neuron differentiation
autonomic nervous system development
forebrain generation of neurons
regulation of bone mineralization
regulation of biomineral tissue development
bone mineralization
regulation of ossification
central nervous system neuron differentiation
biomineral tissue development
neural precursor cell proliferation
mesenchymal cell differentiation
stem cell differentiation
glial cell differentiation
mesenchyme development
forebrain development
pattern specification process
p-value
Fold Enrichment

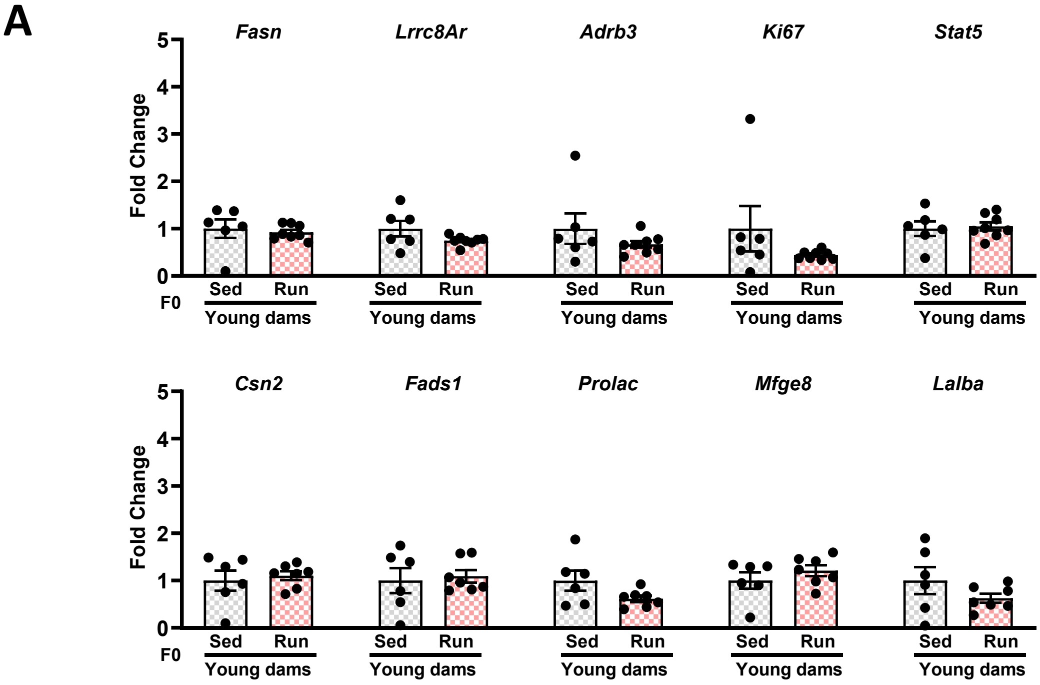
A
Fasn
Lrrc8Ar
Adrb3
Ki67
Stat5
Fold Change
5
4
3
2
1
0
Sed
Run
F0
Young dams
Csn2
Fads1
Prolac
Mfge8
Lalba
